# Thermodynamic Integration in 3n Dimensions without Biases or Alchemy for Protein Interactions

**DOI:** 10.1101/150870

**Authors:** Liao Y Chen

**Affiliations:** Department of Physics, University of Texas at San Antonio, San Antonio, Texas 78249 U.S.A.

## Abstract

Thermodynamic integration (TI), a powerful formalism for computing the Gibbs free energy, has been implemented for many biophysical processes characterized by one-dimensional order parameters with alchemical schemes that require delicate human efforts to choose/design biasing potentials for sampling the desired biophysical events and to remove their artifactitious consequences afterwards. Theoretically, an alchemical scheme is exact but practically, it causes error amplification. Small relative errors in the interaction parameters can be amplified many times in their propagation into the computed free energy [due to subtraction of similar numbers such as (105 ± 5) − (100 ± 5) = 5 ± 7], which would render the results significantly less accurate than the input interaction parameters. In this paper, we present an unsophisticated implementation of TI in 3n dimensions (3nD) (n=1,2,3…) without alchemy or biasing potentials. In TI3nD, the errors in the interaction parameters will not be amplified and human efforts are not required to design biasing potentials that generate unphysical consequences. Using TI3nD, we computed the standard free energies of three protein complexes: trometamol in Salmonella effector SpvD (n=1), biotin in avidin (n=2), and Colicin E9 endonuclease with cognate immunity protein Im9 (n=3) and the hydration energies of ten biologically relevant compounds (n=1 for water, acetamide, urea, glycerol, trometamol, ammonium and n=2 for erythritol, 1,3-propanediol, xylitol, biotin). The computed results all agree with available experimental data. Each of the 13 computations is accomplishable within two (for a hydration problem) to ten (for the protein-recognition problem) days on an inexpensive workstation (two Xeon E5-2665 2.4GHz CPUs and one nVidia P5000 GPU).

## INTRODUCTION

Accurate computation of protein interactions is fundamental to understanding essential biological processes such as molecular recognitions in terms of “the gigglings and wigglings” of the atoms that constitute the biomolecules and their aqueous environments.[1] However, errors are inevitable in all computations including quantification of protein interactions in terms of the Gibbs free energy. Errors in the input force fields parameters propagate to the final results in the free-energy differences we are interested to evaluate. And the relative errors can be amplified catastrophically in the process. Take the hydration of a bio-relevant molecule, the simplest biomolecular interaction, for an example. The powerful alchemical approach gives the free energy of hydration as the difference, ΔG_hydr_ = ΔG_vac_−ΔG_aq_, between the Gibbs free energy to annihilate the molecule in vacuum, ΔG_vac_, and the same term in water, ΔG_aq_. For a hypothetical (but not atypical) molecule, ΔG_vac_ = 100 ± 5 kcal/mol and ΔG_aq_ = 105 ± 5 kcal/mol each with about a 5% error, then the free energy of hydration ΔG_hydr_ = 5 ± 7 kcal/mol in which the relative error is amplified to 140% if the errors in the two annihilation energies do not happen to cancel each other out. Very high accuracy in the free energies of annihilation is necessary to have a reasonable accuracy in the computed free energy of hydration. Indeed, the alchemical methods have been widely applied with many successes but they have also been found to have their share of producing false positives/negatives.[2] It is still challenging for us to accurately compute the absolute binding free-energies for various protein interactions.[3–22]

Thermodynamic integration (TI)[23] is an exact and powerful formalism that has been widely adapted and used in the current literature. Typically, a numerical implementation of TI is to compute the potential of mean force (PMF) [23–27] by integrating the mean force acting on a collective degree of freedom represented with a one-dimensional (1D) order parameter[28]. It may involve alchemical cycles that cause amplifications of the inevitable errors in the force field parameters. And it depends on delicately-designed biasing potentials to achieve sufficient sampling for convergence, which bring in artifactitious effects that must be carefully removed to reach the physics of the natural systems of our investigation. The application of biasing potentials and the subsequent removal of their unphysical consequences may also cause error amplifications from subtractions of similar numbers.

In this paper, we present a direct, unsophisticated implementation of TI in 3n-dimensions (TI3nD), without invoking alchemy or biasing potentials, for noncovalent interactions between proteins, ligands, and aqueous environments. In contrast to the orthodox implementations of TI for 1D PMF as a function of an order parameter representing one degree of freedom of the system which is biased delicately with various biasing potentials, this simple implementation of TI is for 3nD PMF as a function of 3n coordinates of n centers directly representing 3n degrees of freedom of the system without invoking any biasing potentials. The n centers can simply be n atoms of the binding partners, ligands or proteins, forming a binding complex. They can also be n centers-of-mass of n groups of atoms chosen on the binding partners. We tested TI3nD on computing the hydration energies of ten biologically relevant small molecules for which we obtained results in close agreements with the experimental data available from the current literature. We also validated TI3nD by computing the absolute binding free energies of three protein binding problems: (1) Colicin E9 endonuclease (E9 DNase) in complex with the cognate immunity protein Im9[29] gives the most difficult test of TI3nD (n=3) as any protein recognition problems will do. For the E9-Im9 problem that has been well-studied experimentally[30], the absolute binding free energy is yet to be computed on a rigorous basis in the current literature. Computations have only been carried out on the relative binding free energies among various E9 alanine mutants with no definitive agreements with the in *vitro* data[31], which well reflects the problem of error amplifications inherent in alchemical algorithms involving subtractions between similarly large numbers. In this difficult test, three rounds of TI3nD were conducted to achieve convergence in the computed absolute binding free energy. The converged results agree with the in *vitro* data with chemical accuracy. (2) Biotin (BTN) in complex with avidin (AVD) represents the strongest noncovalent binding among ligand-protein interactions, for which the test run of TI3nD (n=2) produced perfect agreement with the in *vitro* data and confirmed the hypothesized importance of the loop connecting the 3^rd^ and the 4^th^ β-sheets of AVD[11]. (3) Trometamol (TRS) in complex with Salmonella enterica effector protein SpvD provided us the simplest case (n=1) in which the computed binding affinity well agrees with the value extracted from the in *vitro* experiments.[32]

## METHODS

### Absolute binding free energy from PMF in 3nD

Following the standard literature[3, 15], through the derivation steps detailed in Refs. [33, 34], one can relate the standard (absolute) free energy of binding to the PMF difference and the two partial partitions as follows:

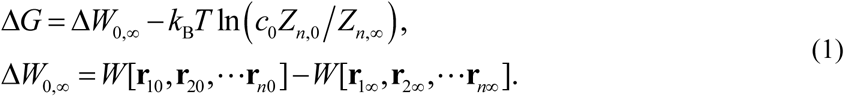

Here *c_0_* is the standard concentration. *k_B_* is the Boltzmann constant. *T* is the absolute temperature. 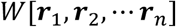 is the 3n-dimensional PMF, the free energy of the system as a function of 3n coordinates of the n centers 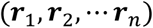 chosen to represent the positions of the partner molecules of the binding complex, which can simply be n atoms or the centers-of-mass of n groups of atoms. The subscripts 0 and ∞ indicate the bound and the dissociated states respectively. Δ*W*_0_,_∞_ is the PMF difference between one state out of the bound state ensemble 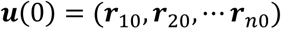 and the corresponding one state of the dissociated state ensemble 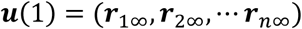. Those two particular states are connected *via* a chosen curve in 3nD space,

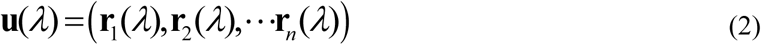

where the parameter *λ* goes from 0 to 1. Along the curve, all the 3n degrees of freedom of the n centers are not allowed to fluctuate. Their fluctuations in the bound state ensemble and in the dissociated state ensemble are represented in the partial partitions *Z_n_*,_0_ and *Z_n_*_,∞_ respectively. The bound state partial partition *Z_n_*_,0_ integrates all 3n degrees of freedom in the neighborhood of the initial state ***u***(0),

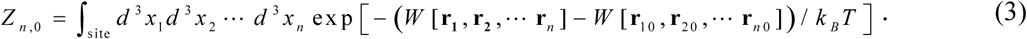

The partial partition function *Z_n_*,_∞_ of the dissociated state has the integration over 3(n–1) degrees of freedom while 3 degrees of freedom are fixed so that the fluctuations in the unbound state would not bring the system back into the bound state,

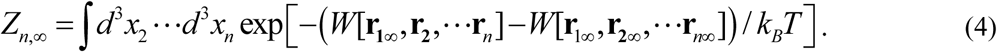

The formulas for computing these partial partitions are given in the supplemental material (SM). With Eqs. (1), (3) and (4), the computations of the fluctuations are isolated in the bound or the unbound state. Along the dissociation path, Eq. (2), the system is free from any fluctuations in all the 3n degrees of freedom. This isolation-of-fluctuations approach reduces computing efforts and enhances accuracy as demonstrated in the three binding problems and ten hydration problems discussed in the Results and Discussion section.

### PMF in 3nD

The central to this work is the PMF in 3nD, 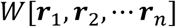, as a function of the 3n coordinates of n centers 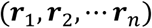 along a single curve/line 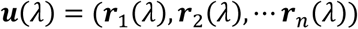 connecting one state 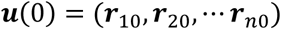 that is arbitrarily chosen from the bound state ensemble to the corresponding one state 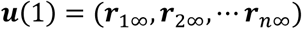that belongs to the dissociated state ensemble. From the definition of PMF, we have the following formula for the 3nD PMF difference:

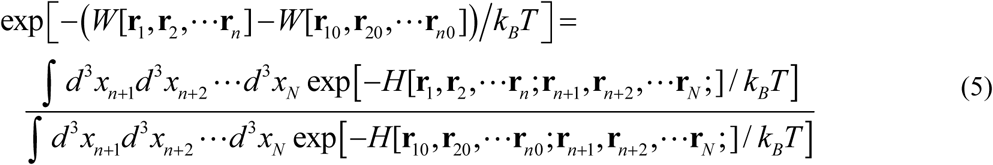

where *H* is the Hamiltonian of the entire system, a function of 3*N* coordinates 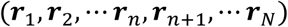 of all the *N* atoms of a model system. A straightforward extension of the well-known TI gives the following formula of TI3nD:

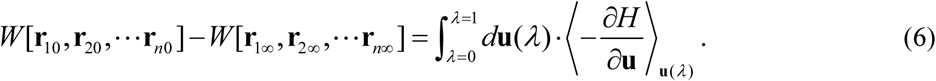

Here the partial derivative with respect to ***u*** means 3nD gradient of the Hamiltonian. The dot product is between the two 3nD vectors. The brackets with subscript ***u***(*λ*) mean the statistical average of the total forces acting on the n centers by all the 3(*N-n*) degrees of freedoms of the system when the n centers are fixed at 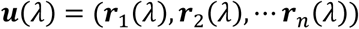. In this, the 3nD PMF difference between two states is equal to the line integral of the mean force acting the 3n degrees of freedom along a line connecting the two states. Since PMF is a function of state, any one line/curve connecting the two end states is necessary and sufficient for the computation in Eq. (6). This TI3nD formula can be implemented without biasing potentials and therefore no efforts are needed to remove the artifactitious effects of the biases.

### Hydration energy from PMF in 3nD

Hydration of a molecule can be regarded as a simple binding problem for which the PMF is along a straight path from deep inside a box of water to far outside the box. In each of the 10 hydration problems studied in this work, the solute molecule was inside a cubic box of water with each side at 80 Å. The coordinates were chosen so that the top side of the cube is parallel to the xy-plane at *z*_0_ = 10Å and molecule is located at the origin, *z_A_* = 20Å. Bring the solute along the z-axis from *z_A_* = 0Å to *z_B_ =* 20Å, the PMF difference Δ*W_A,B_* was computed via Eq. (6). The PMF difference from *z_B_* = 20Å to *z_∞_* = ∞ was computed analytically by approximating water as a continuous medium. For this range, the attraction on the charged solute (net charge q) by the water box can be accurately approximated by the force from its image charge *q*′ = −*q*(*ε/ε*_0_ − 1)/(*ε*/*ε*_0_ + 1)[35]. Consequently,

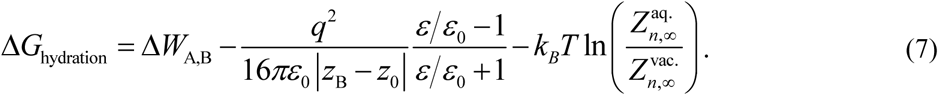

Here *ε*_0_ is the dielectric constant of vacuum. *ε =* 81*ε*_0_ is the dielectric constant of water. Δ*W_A,B_* is the 3nD PMF difference between the state of the molecule being inside water and the state of the molecule being in vacuum. The two partial partitions 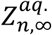 and 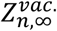 represent the fluctuations in all the other degrees of freedom of a molecule when n centers are fixed in water and in vacuum respectively. For short molecules, one center is sufficient to represent the molecule’s position, the choice of n=1 suffices and 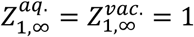. For long molecules, the choice of n=2 can significantly reduce the computing time needed in a given problem because two centers on a molecule can define its position and, partly, its orientation. The tradeoff is that one now must compute the partial partitions in vacuum and in water, as described in SM, Eq. (S5).

### Simulation parameters

In all the MD runs, the CHARMM36 force field parameters[36, 37] were used for all the intra- and inter-molecular interactions. The Langevin stochastic dynamics was implemented with NAMD[38] to simulate the systems at constant temperature of 298 K and constant pressure of 1 bar. The time step was 1 fs for the short-range and 2 fs for the long-range interactions. The damping constant was 5/ps. Explicit solvent (water) was represented with the TIP3P model. Full electrostatics was implemented *via* particle mesh Ewald at the level of 128×128×128 for the three binding problems or *via* the analytical approximation in Eq.(7) for the hydration problems.

### All-atom model systems

The 13 model systems were set up as follows:

1. The Im9-E9 system was formed by taking Chains A and B of the Im9-E9 DNase complex (PDB: 1EMV), translating it to center around the origin of the coordinate systems, rotating it so that the dissociation path is approximately along the z-axis, solvating it with an 80Å×80Å×120Å box of water, neutralizing the its charge with Na^+^/Cl^−^ ions and salinizing it to 150 mM of NaCl.
2. The BTN-AVD system was formed by taking Chain A of the BTN-AVD complex (PDB: 2AVI) and following the same procedure stated in (1).
3. The TRS-SpvD system was formed by taking the X-ray structure of the TRS-SpvD complex (PDB: 5LQ7) and following the procedure described in (1).
4. The hydration system of a small solute (water, acetamide, urea, glycerol, trometamol or ammonium) was formed by placing the solute at the origin of the coordinate system and solvating it with an 80Å×80Å×80Å box of water. The top side of the water box is the plane of z = 10Å in parallel to the xy-plane. The bottom side of the water box is the plane of z = −70Å
5. The hydration system of a long solute (1,3-propanediol, erythritol, xylitol or biotin) was formed by rotating the solute so that it lies on the xy-plane and solvating it in the same way as described in (4).

## RESULTS AND DISCUSSION

The binding free energies of three protein complexes are tabulated in Table I. The computing costs were approximately four days for a drug-protein binding problem (TRS-SpvD or Biotin-Avidin) and ten days for a protein recognition problem (E9 DNase-Im9) on a workstation with dual 8-core CPUs and single nVidia P5000 GPU. The hydration energies of ten small molecules are tabulated in Table II. The computing cost for a hydration problem is less than two days on the same workstation. All these computations do not amplify the inherent errors of the force field parameters, namely, the errors in the final results remained around the level of *±k_B_T.* And it is worth noting that the TI3nD computing cost for an absolute binding energy is similar to the state-of-the-art approaches for relative binding energies of current literature, *e.g*., Ref. [39].

**Table I.** Experimental vs computed biding free energies

| Binding complex | PDB code | Experimental $\Delta G$ (kcal/mol) | This work $\Delta G$ (kcal/mol) | $\Delta W_{0,\infty}$ (kcal/mol) | $k_B T \ln \left( \frac{Z_{n,\infty}}{c_0 Z_{n,0}} \right)$ (kcal/mol) | Simulation time (ns) |
| --- | --- | --- | --- | --- | --- | --- |
| Im9-E9DNase | 1EMV | -18.6[29] | -19.3 | -39.8 | 20.5 | 450* |
| Biotin-Avidin | 2AVI | -21.0[40] | -20.5 | -29.8 | 9.3 | 175* |
| TRS-SpVD | 5LQ7 | -4.5[32] | -4.1 | -9.5 | 5.4 | 165* |

\*These times do not include the extra rounds of simulations conducted to demonstrate convergence of the computations.

**Table II.**
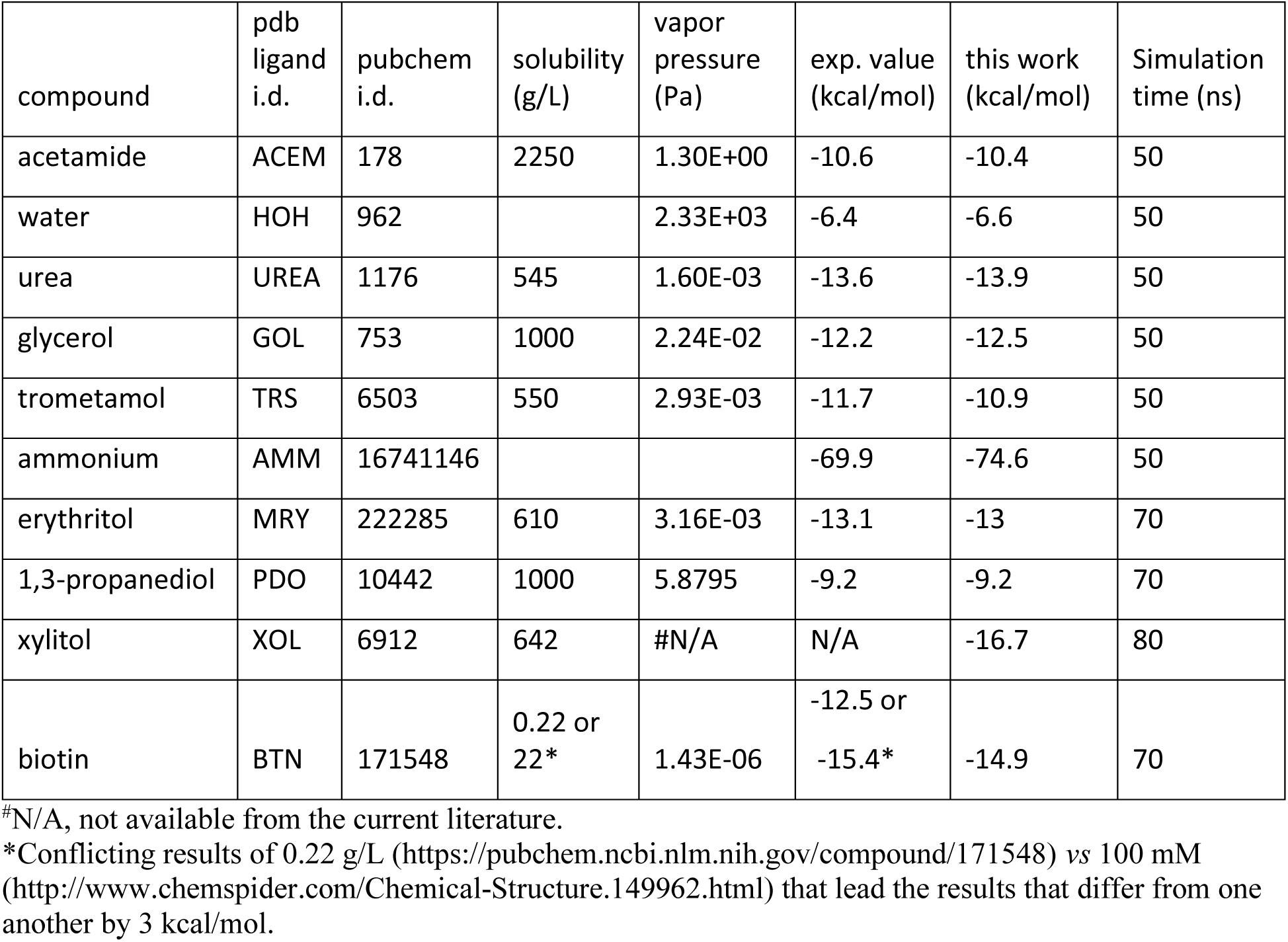
Experimental vs. computed hydration energies

### Im9-E9 DNase

Colicin E9 endonuclease (E9 DNase) binds very strongly with its cognate immunity protein Im9. This binding complex has received intensive investigations for its direct relevance in bacterial biology and for its being a model system for the study of how one protein recognizes another[29–31, 41]. This difficult binding problem represents a serious challenge for TI3nD or any other theoretical-computational approaches to produce quantitative agreement with the in *vitro* data available in the current literature. To meet this standing challenge, we conducted multiple runs of atomistic MD simulations based on the established CHARMM 36 force field parameters[36] from which we obtained the absolute free energy of Im9-E9 binding in quantitative agreement with the in *vitro* data (Table I). This close agreement between the in *silico* and the in *vitro* data indicates that the chemical accuracy (*±k_B_T*) can be achieved for protein recognition problems when rigorous statistical mechanics is implemented on the basis of the established force field parameters. In the implementation of TI3nD for Im9-E9, three α-carbons were chosen on the immunity protein Im9 (Asn24Cα, Leu33Cα, and Tyr54Cα) as the three centers whose coordinates represent the position and orientation of Im9 (Fig. 1, Inset). The PMF shown in Fig. 1 is a function of the displacement of these three centers (which are displaced identically). The displacement from 0 to 6 Å was divided into 60 increments (or windows) of 0.1 Å each and the displacement from 6 Å to 12 Å was divided into 30 increments of 0.2 Å each. At each of these 90 displaced positions of Im9, four sets of data of the forces acting the three centers were collected from four segments of unbiased MD runs after a long equilibration process during which the three centers on Im9 and, additionally, the 22 centers on E9 (detailed later in this subsection) were fixed. The statistical mean of the forces was integrated as in Eq. (6) along the displacement to yield the PMF shown in Fig. 1.

**Fig. 1.**
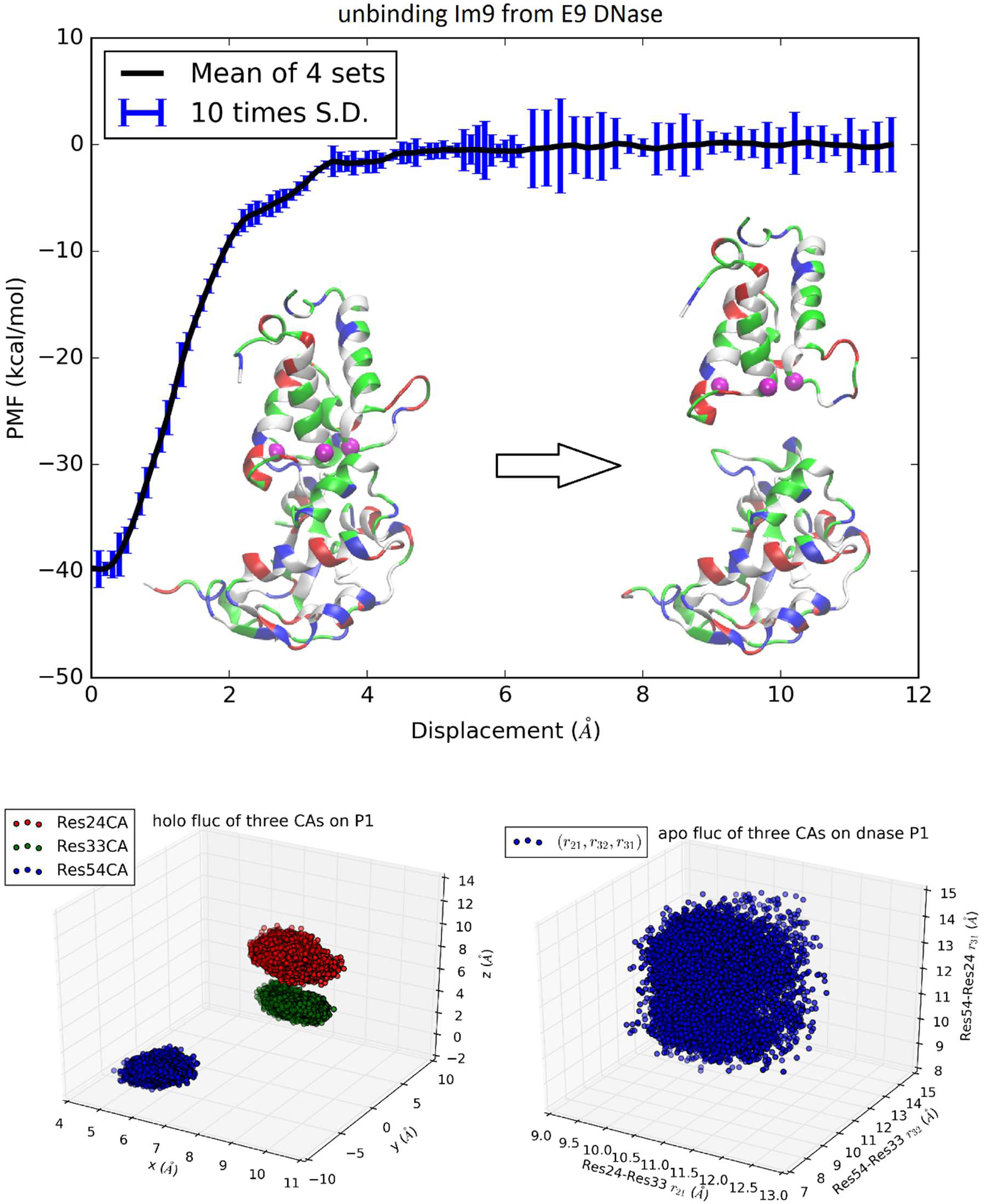
Unbinding Im9 immunity protein form Colicin E9 DNase. Shown in the top panel is the PMF curve of unbinding Im9 from E9. Insets: Im9 and E9 in the bound state (left) and in the unbound state (right) are shown in ribbons colored by residue types (hydrophobic, white; hydrophilic, green; negatively charged, red; positively charged, blue). The three centers (α-carbons) on Im9 (Asn24Cα, Leu33Cα, and Tyr54Cα) are marked as purple balls, whose coordinates represent the position and orientation of Im9. The bottom left panel shows the fluctuations of the three centers in the bound state in terms of their coordinates: (*x*_1_, *y*_1_, *z*_1_), (*x*_2_,*y*_2_, *z*_2_), and (*x*_3_, *y*_3_*,z*_3_). The bottom right panel shows their fluctuations in the unbound state in terms of three mutual distances: (*r*_21_, *r*_32_*, r*_31_).

The convergence behavior of TI3nD for PMF is illustrated in Fig. 2 that shows the results of three runs of simulations. In Run 1, the long equilibration was 1 ns in duration and the force-collecting segments were 0.1 ns each in duration. In Run 2, the long equilibration was 5.4 ns in duration and the force-collecting segments were 1 ns each in duration. In Run 3, the long equilibration was 9.4 ns in duration and the force collecting segments were 1 ns each in duration. Fig. 2 indicates that Run 2 was needed to produce converged results in (0, 6 Å) and the Run 1 was needed in (6 Å, 12 Å). Therefore, the necessary computing cost (Table I) is slightly less than half of the computations we carried out to demonstrate the convergence.

**Fig. 2.**
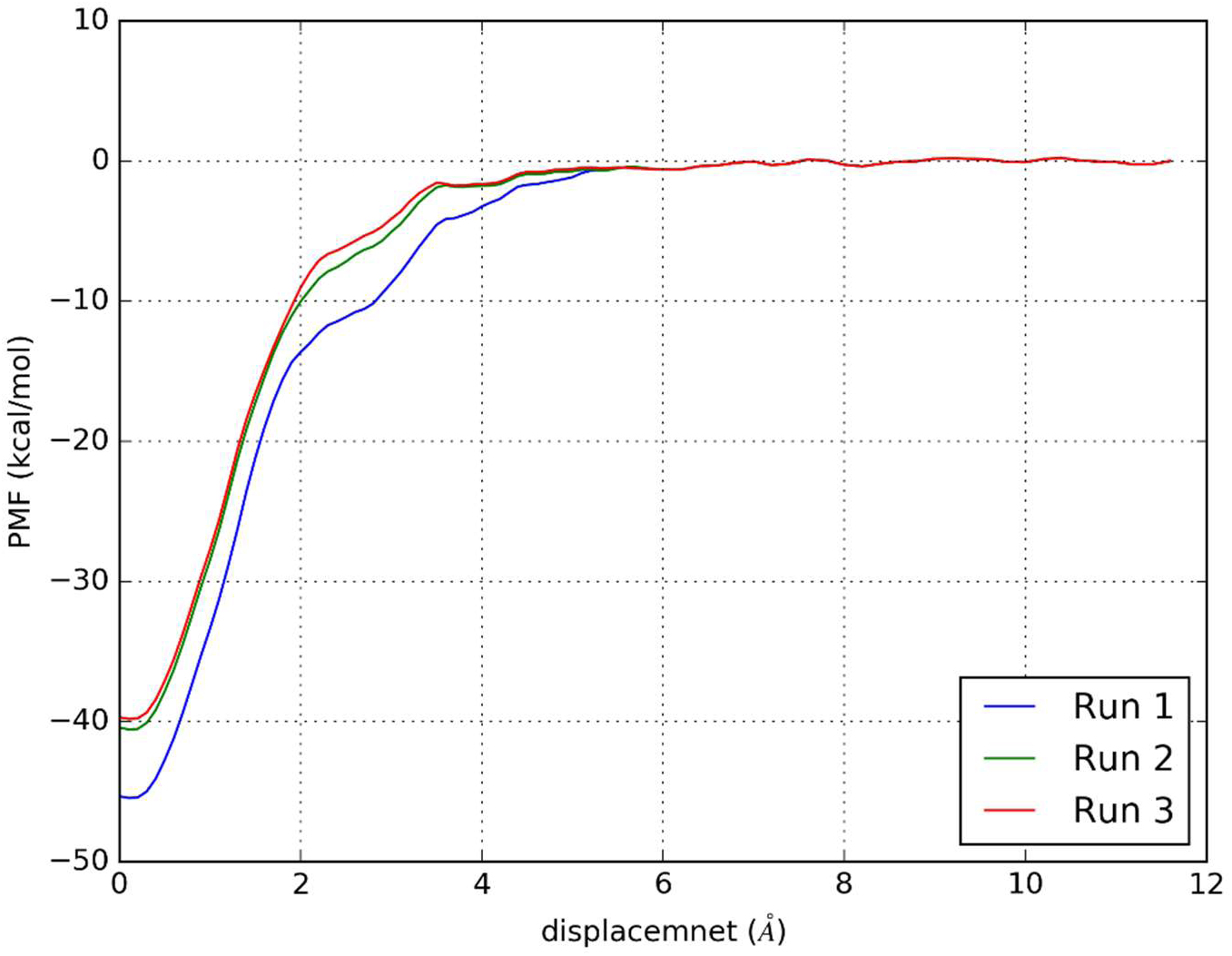
Convergence of TI3nD approach for Im9-E9 DNase binding free energy calculation. The curves represent the PMF as a function of Im9 displacement from the binding site in three runs. There are 60 windows of 0.1 Å each in width from 0 to 6.0 Å and 30 windows of 0.2 Å each in width from 6.0 Å to 12 Å. Run 1, the PMF gradients were the statistical mean of four sets of data collected from four 0.1 ns simulations after 1 ns equilibration in each window. Run 2, the PMF gradients were the statistical mean of four sets of data collected from four 1 ns simulations after 1.4 ns equilibration in each window. Run 3, the PMF gradients were the statistical mean of four sets of data collected from four 1 ns simulations after 5.4 ns equilibration in each window. The deviations between Run 2 and Run 3 are within the margin of error, *±k_B_T.*

In order to account for the E9 conformation-rigidity changes due to binding, 22 α-carbons on E9 DNase (Pro73Cα, Ser74Cα, Asn75Cα, Lys76Cα, Ser77Cα, Ser78Cα, Val79Cα, Ser80Cα, Phe86Cα, Lys89Cα, Asn90Cα, Gln91Cα, Val93Cα, Arg96Cα, Val98Cα, Pro124Cα, Lys125Cα, Arg126Cα, His127Cα, Ile128Cα, Asp129Cα, and Ile130Cα) were chosen as the 22 centers for which we computed the changes in deviations and fluctuations between the bound and the unbound states. These 22 centers were fixed during the simulations covering the entire range of Im9 displacements. Therefore, n=3+22 in this case of TI3nD study. The two partial partitions involved in the binding energy formula, Eq. (1), are 75-dimensional and 72-dimensional, respectively. Namely, *Z*_*n*,0_ = 2.1×10^−22^Å^75^ and *Z_n_*,_∞_ *=* 1.1×10^−10^Å^72^. The ratio between these two partial partitions constitutes a large part of the absolute binding energy in Eq. (1) (Table I). The contributors to these two partial partitions are the following:

The position and orientation of Im9 are quantified by the nine coordinates of the three centers on Im9 (Asn24Cα, Leu33Cα, and Tyr54Cα shown in insets of Fig. 1). The nine degrees of freedom of these three centers can be divided into the first group of six degrees of freedom for the translation and rotation of Im9 and the second group of three degrees of freedom for its rigidity. In the bound state, the first group of six degrees of freedom are confined by the binding partner E9 (Fig. 1, bottom left panel). In the unbound state, they are unconfined, namely, Im9 is free to translate (three degrees of freedom) and to rotate (three degrees of freedom). The confinement of the first group of six degrees of freedom corresponds to a contribution of 13.77 kcal/mol to the TI3nD computation of the binding free energy. The second group of three degrees of freedom for the relative motions between these three centers (represented by the three mutual distances shown in Fig. 1, bottom right panel) does not differ significantly between the bound and the unbound states. The relative shifting of these three centers contributes −0.98 kcal/mol and the rigidifying of them upon binding, 1.35 kcal/mol, to the absolute binding energy, which sum up to 0.37 kcal/mol.

The position and situation of E9 are quantified by the coordinates of the 22 centers chosen on/near the contact surface with Im9. Upon binding with Im9, they become slightly more rigid. The reduction of their fluctuations altogether give rise to a contribution of 2.23 kcal/mol to the absolute binding energy. The shifting of these 22 centers contributes 5.13 kcal/mol to the absolute binding energy. Both factors are positive and thus against E9-Im9 binding. The factors for Im9-E9 binding, which give rise to the steep changes in the PMF curve of Fig. 1, are the hydrophobic contacts and the buried hydrogen bonds between the two proteins (SM, Fig. S1).

### BTN-AVD

The binding problem of the biotin-avidin complex has been a subject of repeated theoretical-computational investigations because it represents the strongest non-covalent binding and has brought serious challenges to theoretical-computational methods.[11] In fact, optimization of BTN charge distribution led to significant difference from the generic CHARMM values that are shown in SM, Table S1 and Fig. S1. (BTN has a total charge of −*e* in this work at neutral pH.) With the optimized parameters, we obtained the hydration energy of BTN in reasonable approximation to the experimental data (detailed in the next subsection) and the BTN-AVD binding free energy in perfect agreement with the experimental data (Table I). Since BTN is a long molecule, it is appropriate to choose n=2. Atoms C4 and C10 of BTN (SM, Fig. S1) are chosen as the two centers (six degrees of freedom) to represent its location and orientation. Along one dissociation path, the PMF was computed as integration of the mean forces on the six degrees of freedom in dot product with the 6D displacement as defined in Eq. (6). Four sets of force data were collected from four segments of 0.1 ns sampling after 1 ns equilibrium at each given position. The mean values and the standard deviations are shown Fig. 3, top panel. The middle panel of Fig. 3 shows the fluctuations of BTN-C4-C10 in the bound state ensemble, which gives the bound state partial partition *Z*_2,0_ = 0.209Å^6^. The bottom panel of Fig. 3 shows the fluctuations of BTN-C4-C10 in the unbound state ensemble in terms of one degree of freedom that is the C4-C10 distance fluctuating in time.

**Fig. 3.**
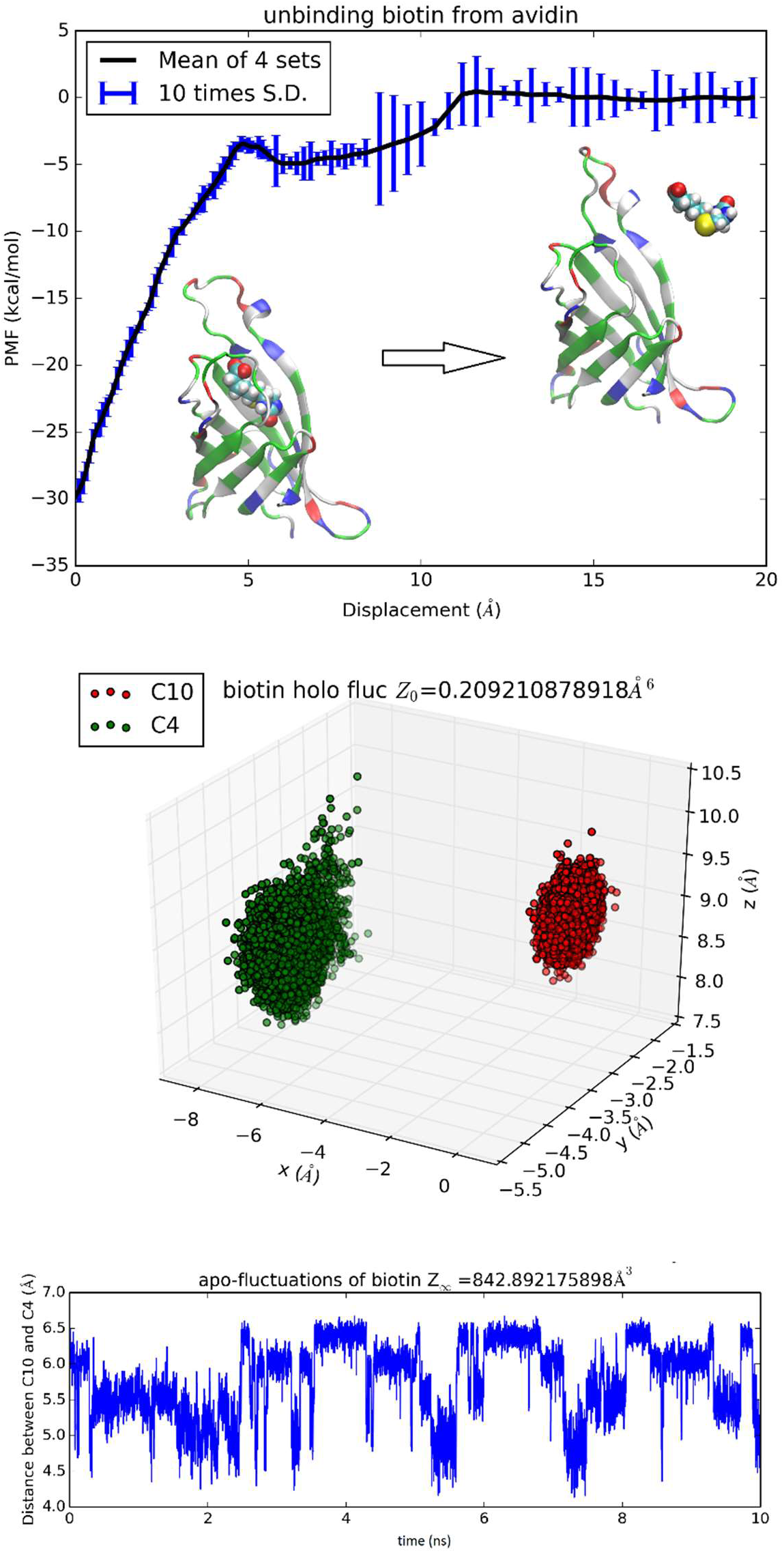
Unbinding biotin from avidin. In the top panel, the PMF is plotted as a function of the displacement of BTN along an unbinding path. Inset: the protein is shown as ribbons colored by residue types and biotin is shown as spheres colored by atoms (hydrogen, white; carbon, cyan; oxygen, red; nitrogen, blue; sulfur, yellow). Fluctuations of C4 and C10 in the bound state ensemble are shown in the middle panel. The fluctuating distance between C4 and C10 in the unbound state ensemble is shown in the bottom panel.

In the unbound state ensemble, two degree of freedom are in rotation where BTN is free to rotate and its environment is isotropic. These one plus two degrees of freedom in the unbound state combine to give the unbound state partial partition *Z*_2_,_∞_ = 842.9Å^3^. From the computed values of the PMF difference and the two partial partitions, we obtained the absolute binding free energy of biotin-avidin Δ*G* = −20.5kcal/mol which is in perfect agreement with the experimental value of −21.0kcal/mol.

It is interesting to examine the interactions between BTN and the loop connecting the 3^rd^ and the 4^th^ β-sheets consisting of residues 35 to 46 (noted as L3,4) which were suggested as a significant contributor to the overall binding free energy.[11] The van der Waals interaction energy between the charged BTN and L3,4 is shown in Fig. 4 (central panel) as a function of BTN displacement from the binding packet (left panel) along the dissociation path to the unbound state (right panel). With a signal clearly above the noise level, this curve indicates a contribution of about −6 kcal/mol to the total binding free energy of −20.5 kcal/mol, which confirms the hypothesis of Ref. [11]. The electrostatic energy between BTN and L3,4 is also shown in Fig. 4. However, the noise is not significantly lower than the signal in this case and thus prevents us from drawing any conclusions based on this one curve. Besides, factors including L3,4-water and BTN-water interactions need to be incorporated and the cancellations among them will cause gross amplifications of the relative errors in approaches relying on direct computation of energies. These considerations all favor the PMF-based approaches for quantitative accuracy.

**Fig. 4.**
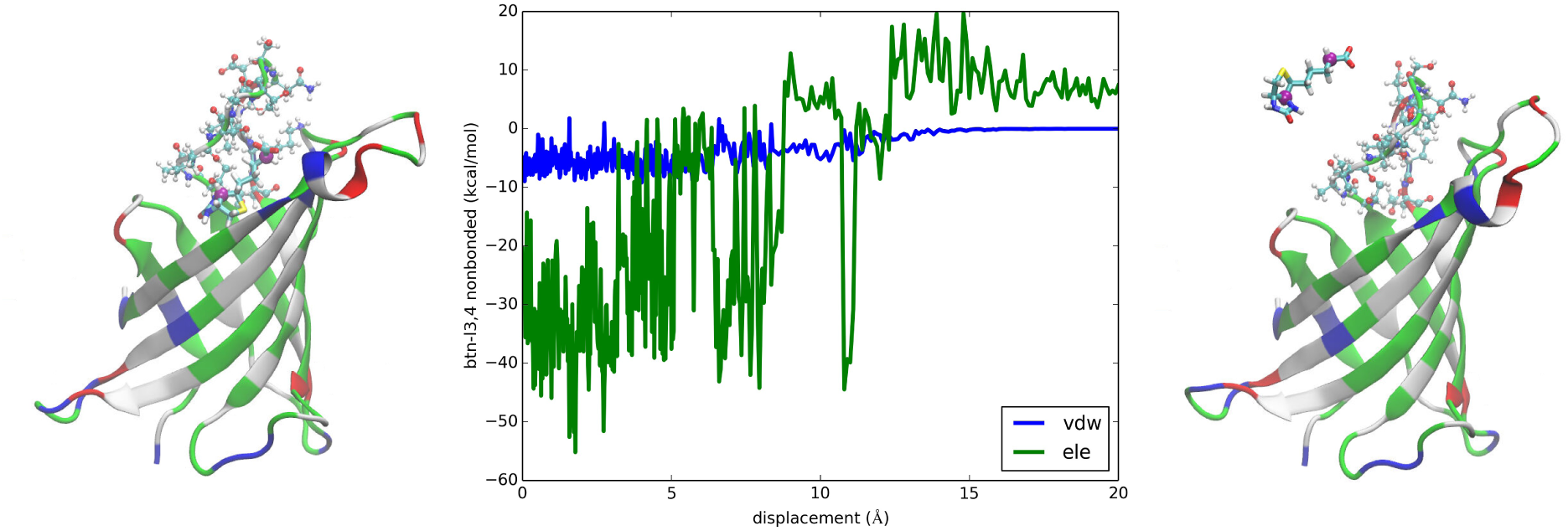
Interactions between biotin and L3,4 of avidin. Plotted in the central panel are the van der Waals and the electrostatic interactions between biotin and L3,4 along the dissociation path. The left and the right panels show the bound and the unbound states of BTN-AVD, respectively. The whole protein is shown as ribbons colored by residue types. The L3,4 residues are shown in ball-and-sticks colored by atom names. BTN is shown in licorices colored by atom names whose C4 and C10 atoms are shown as purple spheres.

### TRS-SpvD

SpvD is a Salmonella enterica effector protein whose structure was resolved to atomistic resolution to elucidate the mechanistic basis of its role as a cysteine protease.[32] The binding of a trometamol (TRS) in the structure was not identified as biologically relevant, which were due to the fact that 100 mM of TRS was present in the buffer liquids. Nonetheless, this binding complex provides a simple test for the TI3nD technique because its absolute free energy of binding can be readily inferred from the experimentally measured values of its occupancy at the binding site (PDB code: 5LQ7). In this test of TI3nD, we chose n=1. The tetrahedral carbon of trometamol (TRS-C) was chosen as the center to the represent the molecule’s location (three degrees of freedom). Along one dissociation path in the 3D space of the TRS-C coordinates, the PMF was computed as the integration of the mean forces on the three degrees of freedom in dot product with the 3D displacement as defined in Eq. (6). Four sets of force data were collected from four segments of 0.1 ns sampling after 1 ns equilibrium at each given position as indicated by the error bars in Fig. 5. The mean values and the standard deviations are shown Fig. 5, top panel. Fig. 5 also shows the 3D fluctuations of TRS-C at the binding site that gives rise to the partial partition of the bound state *Z*_1,0_ = 0.198Å^3^. In this case of n=1, the partial partition of the unbound state is simply *Z*_1,∞_ = 1. The PMF difference between the bound and the unbound states Δ*W*_0,∞_ = −9.5kcal/mol and two partial partitions combine to give the absolute binding free energy Δ*G* = −4.1kcal/mol which well agrees with the experimental data of −4.5 kcal/mol.

**Fig. 5.**
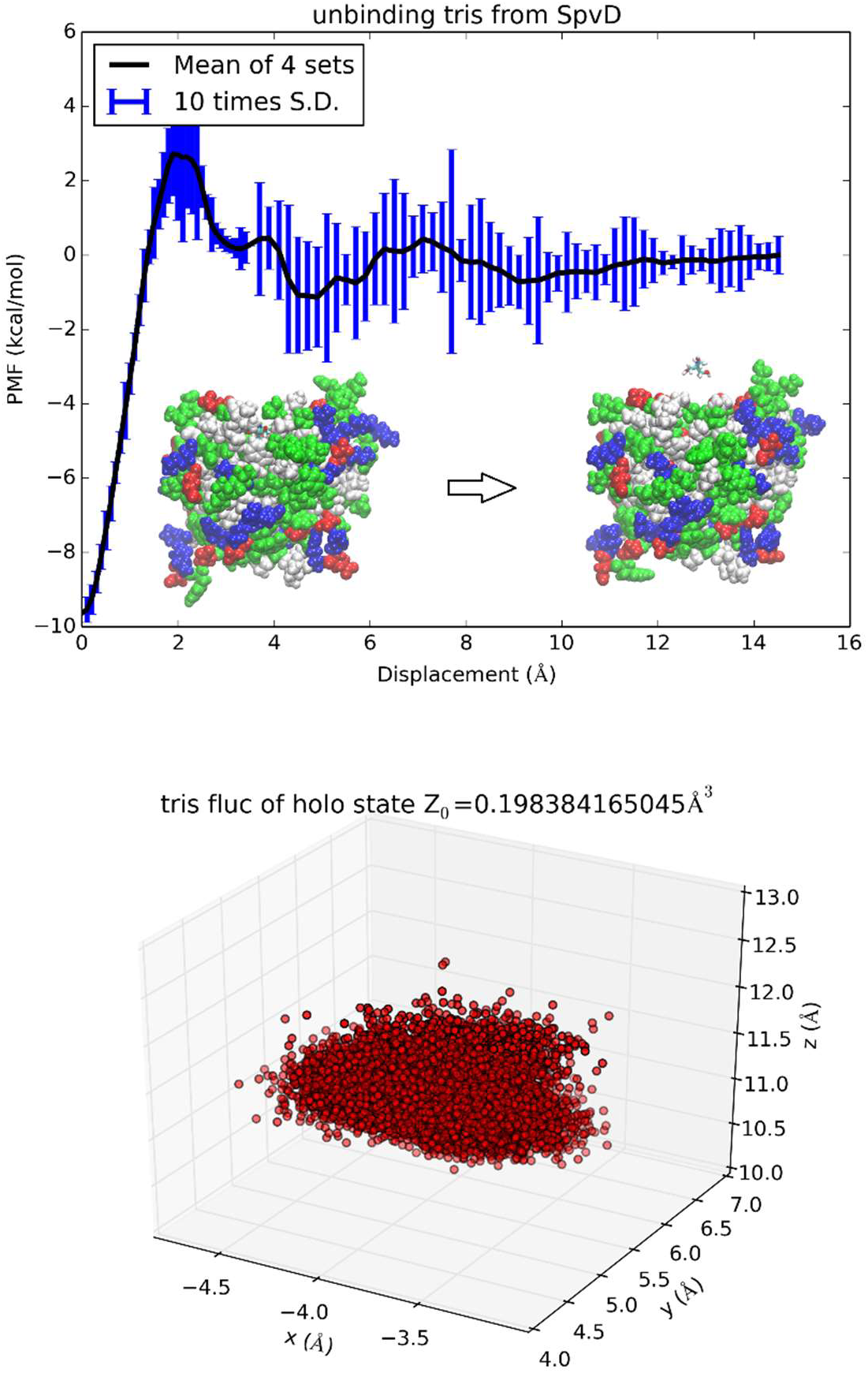
Unbinding trometamol from Salmonella SpvD. In the top panel the PMF is shown as function of the displacement of the TRS tetrahedral carbon (TRS-C) along the z-axis. Inset: the protein is shown as spheres colored by residue types and TRS is shown as licorices colored by atoms. The bottom panel shows TRS fluctuations at the binding site (the xyz-coordinates of TRS-C sampled in the bound state ensemble).

### Hydration energy

For acetamide, water, urea, glycerol, trometamol and ammonium, one center (n=1) was chosen to represent the molecule’s position and therefore partial partitions in water and in vacuum are both equal to one, 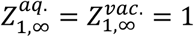. The third term on the right-hand side of Eq. (7) is simply zero. The PMF difference (shown in SM, Figs. S3 to S8) gives directly the hydration free energy except the case of ammonium in which the net charge of the cation render the second term of Eq. (7) nonzero. For erythritol, 1,3-propanediol, xylitol, and biotin, two centers (n=2) were chosen to represent the molecule’s position. The partial partitions in water and in vacuum give a small but nonzero contribution in the third term of Eq. (7). In these four cases, 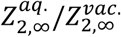 can be computed by invoking the Gaussian approximation for the fluctuations or by integrating the PMF for stretching the molecule between the two centers (SM, Eq. (S5)). The PMF curves and the partial partitions are shown in Fig. 6 for biotin and in SM, Figs. S8 to S11 for erythritol, 1,3-propanediol and xylitol respectively. The computed hydration energies of these ten biologically relevant compounds are tabulated in Table II alongside the experimental data. In each case, the computed free energy of hydration agrees closely with the experimental data available from the current literature.

**Fig. 6.**
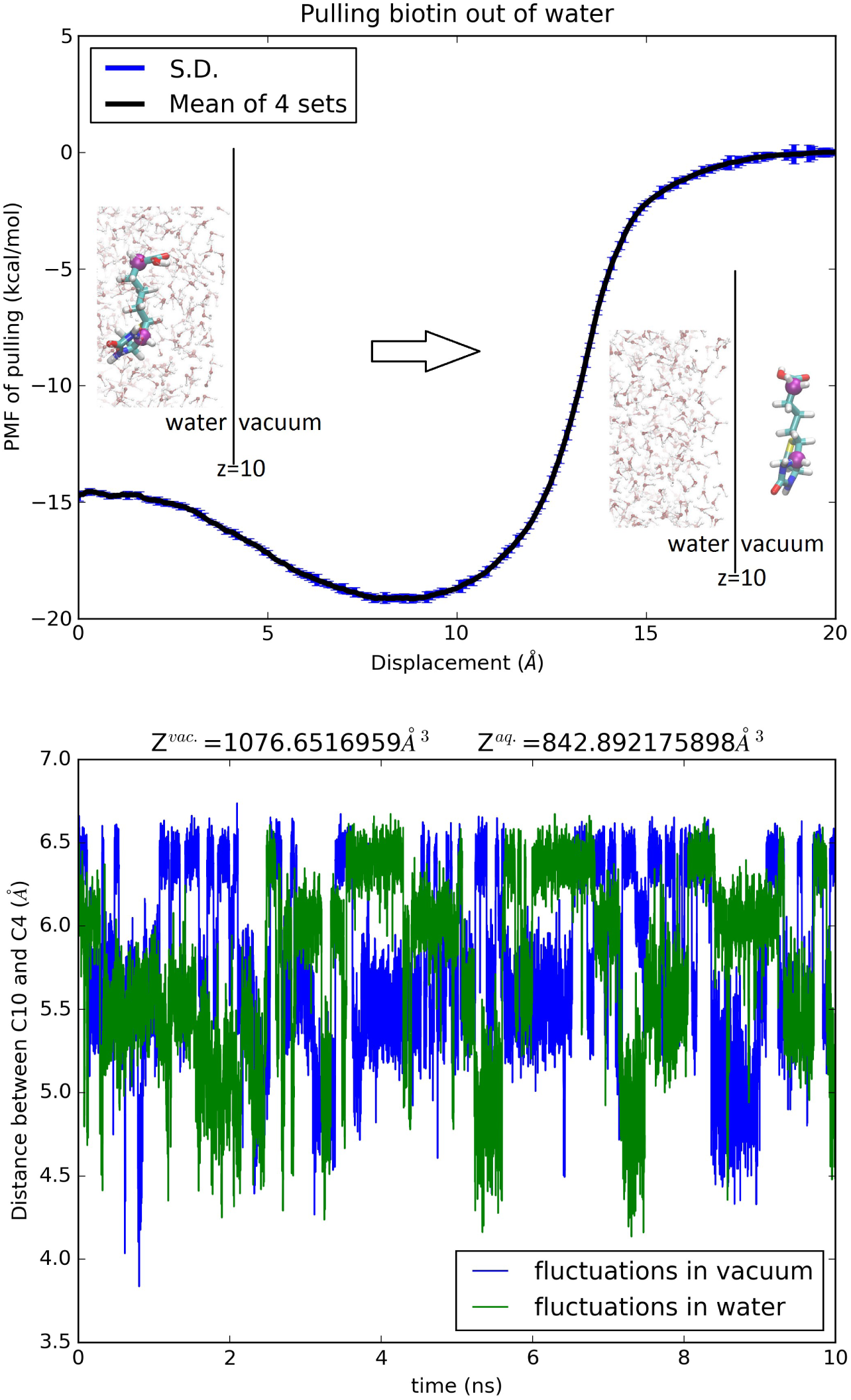
Hydration of biotin. Top panel, the PMF of pulling biotin out of water. The error bars reflect the standard errors among four sets of samplings. The insets show the protonated, neutral biotin in water (left) and outside water (right). BTN is shown as licorices colored by atom names whose C4 and C10 atoms are shown as purple spheres indicating the two centers chosen in this study. The water-vacuum interface is at z=10 Å. Some of the waters are shown in ball-and-sticks colored by atom names. The bottom panel shows the fluctuating distance between C4 and C10 in water and in vacuum.

## CONCLUSIONS

In terms of computational approaches, TI3nD is an unsophisticated implementation of TI on a rigorous basis of statistical mechanics. Without invoking biasing potentials or alchemical schemes, it is shown to be accurate and efficient for protein-interaction problems and to provide a simple, direct way of computing hydration energies. In terms of biological understandings, quantitative insights have been elucidated on the thermodynamic characteristics of E9-Im9 recognition and on the strength of biotin-avidin binding, both in perfect agreements with in *vitro* experiments. With TI3nD, many complex biophysical processes can be investigated with quantitative accuracy at moderate computing costs.

## SUPPLEMENTAL MATERIAL

The online-only supplemental material (SM) contains equations, tables, and figures that are discussed but included in the main context.

## ACKNOWLEDGEMENTS

The author acknowledges support from the NIH (GM121275) and the computing resources provided by the Texas Advanced Computing Center at University of Texas at Austin.

## REFERENCES

[1] J. D. Durrant, and J. A. McCammon, BMC Biol. 9, 71 (2011).

[2] S. E. Boyce et al., J. Mol. Biol. 394, 747 (2009).

[3] H.-J. Woo, and B. Roux, Proc. Natl. Acad. Sci. U. S. A. 102, 6825 (2005).

[4] F. M. Ytreberg, R. H. Swendsen, and D. M. Zuckerman, J. Chem. Phys. 125, 184114 (2006).

[5] D. L. Mobley, and K. A. Dill, Structure 17, 489 (2009).

[6] H.-X. Zhou, and M. K. Gilson, Chem. Rev. 109, 4092 (2009).

[7] T. Hou et al., J. Chem. Inf. Model. 51, 69 (2010).

[8] L. Cai, and H.-X. Zhou, J. Chem. Phys. 134, 105101 (2011).

[9] J. D. Chodera et *al.*, Curr. Opin. Struct. Biol. 21, 150 (2011).

[10] E. Gallicchio, and R. M. Levy, in Advances in Protein Chemistry and Structural Biology, edited by C. Christo (Academic Press, 2011), pp. 27.

[11] I. J. General, R. Dragomirova, and H. Meirovitch, J. Phys. Chem. B 116, 6628 (2012).

[12] X. Wu, A. Damjanovic, and B. R. Brooks, in Adv. Chem. Phys. (John Wiley & Sons, Inc., 2012), pp. 255.

[13] J. C. Gumbart, B. Roux, and C. Chipot, J. Chem. Theory Comput. 9, 794 (2013).

[14] F. Zeller, and M. Zacharias, J. Phys. Chem. B 118, 7467 (2014).

[15] S. Doudou, N. A. Burton, and R. H. Henchman, J. Chem. Theory Comput. 5, 909 (2009).

[16] M. Misini Ignjatović et al., J. Comput.-Aided Mol. Des., 1 (2016).

[17] L. J. Kingsley et al., J. Comput. Chem. 37, 1861 (2016).

[18] Z. Zhang et al., Biophys. Chem. 211, 28 (2016).

[19] G. Hu et al., Scientific Reports 5, 16481 (2015).

[20] M. Musgaard, and P. C. Biggin, J. Chem. Inf. Model. 56, 1787 (2016).

[21] Y. Niu et al., Chemom. Intell. Lab. Syst. 158, 91 (2016).

[22] P. D. Siders, Mol. Simul. 42, 693 (2016).

[23] J. G. Kirkwood, J. Chem. Phys. 3, 300 (1935).

[24] D. Chandler, J. Chem. Phys. 68, 2959 (1978).

[25] L. R. Pratt, G. Hummer, and A. E. Garcia’, Biophys. Chem. 51, 147 (1994).

[26] B. Roux, Comput. Phys. Commun. 91, 275 (1995).

[27] T. W. Allen, O. S. Andersen, and B. Roux, Biophys. Chem. 124, 251 (2006).

[28] E. Darve, in Free Energy Calculations: Theory and Applications in Chemistry and Biology, edited by C. Chipot, and A. Pohorille (Springer Berlin Heidelberg, Berlin, Heidelberg, 2007), pp. 119.

[29] R. Wallis et al., Biochemistry 34, 13751 (1995).

[30] G. Papadakos, J. A. Wojdyla, and C. Kleanthous, Q. Rev. Biophys. 45, 57 (2011).

[31] A. H. Keeble et al., J. Mol. Biol. 379, 745 (2008).

[32] G. J. Grabe et al., J. Biol. Chem. 291, 25853 (2016).

[33] L. Y. Chen, Journal of Chemical Theory and Computation 11, 1928 (2015).

[34] R. A. Rodriguez, L. Yu, and L. Y. Chen, J. Chem. Theory Comput. 11, 4427 (2015).

[35] L. Y. Chen, D. A. Bastien, and H. E. Espejel, Phys. Chem. Chem. Phys. 12, 6579 (2010).

[36] R. B. Best et al., J. Chem. Theory Comput. 8, 3257 (2012).

[37] K. Vanommeslaeghe, and A. D. MacKerell Jr, Biochimica et Biophysica Acta (BBA) - General Subjects 1850, 861 (2015).

[38] J. C. Phillips et al., J. Comput. Chem. 26, 1781 (2005).

[39] L. Wang et al., J. Am. Chem. Soc. 137, 2695 (2015).

[40] O. Livnah et al., Proc. Natl. Acad. Sci. U. S. A. 90, 5076 (1993).

[41] Y. C. Kim, A. W. Tarr, and C. N. Penfold, Biochimica et Biophysica Acta (BBA) - Molecular Cell Research 1843, 1717 (2014).

